# Motor Occupancy Drives Emergent Mechanical States in Cardiac Myosin Ensembles

**DOI:** 10.64898/2026.07.17.739215

**Authors:** Omayma Y. Alazzam, Md. Amzadul Hoque Chowdhury, Heath M. Stevens, Dana N. Reinemann

**Affiliations:** Department of Biomedical Engineering, University of Mississippi, University, MS, USA; Department of Chemical Engineering, University of Mississippi, University, MS, USA

**Keywords:** myosin, actomyosin actin, ensembles, emergent mechanics, optical trapping

## Abstract

Myosin II generates force through the collective action of mechanically coupled motor ensembles, yet the mechanisms by which these ensembles sense changes in motor occupancy and coordinate force generation remain poorly understood. Ensemble force production may be governed by an optimal balance between effective motor occupancy and mechanical coordination rather than by motor number alone. We reconstituted cardiac myosin ensembles and systematically perturbed effective motor occupancy using the small-molecule drugs omecamtiv mecarbil (OM), which prolongs actomyosin interactions, and mavacamten (MAVA), which reduces the number of available force-generating myosin heads. Optical trapping measurements of full-length and S1 cardiac myosin ensembles revealed that force generation depended on both myosin concentration and pharmacological perturbation. Reducing myosin concentration increased force generation in the absence of drug, while OM and MAVA produced responses that varied with the initial occupancy state of the ensemble. Low concentrations of MAVA enhanced force generation under high motor occupancy but reduced force under low motor occupancy, whereas OM produced occupancy-dependent changes in both endpoint force and force dynamics. Force traces further revealed changes in the persistence and temporal coordination of force generation. These findings support a model in which cardiac myosin ensembles operate along an occupancy-coordination landscape, where productive force generation is achieved at an intermediate level of effective motor occupancy. Our results highlight the importance of effective motor occupancy in shaping emergent ensemble mechanics and suggest that force generation reflects a balance between motor availability and mechanical coordination.

**Statement of Significance:** Force generation by muscle emerges from the coordinated activity of myosin ensembles, yet the principles governing this collective behavior remain poorly understood. Using an in vitro force assay with full-length and truncated cardiac myosin, we systematically perturbed ensemble activity by varying myosin availability and pharmacologically altering the fraction of force-generating motors. We find that force production depends on the initial motor occupancy state of the ensemble, with changes in motor availability producing distinct effects on mechanical output. These findings highlight effective motor occupancy as an important regulator of collective force generation and provide insight into how the interactions of individual molecular motors contribute to the mechanics underlying muscle contractility.

## Introduction

Muscle myosin II converts the chemical energy of ATP hydrolysis into mechanical work through cyclic interactions with actin filaments (1–6). Because individual myosin II molecules are non-processive and spend much of their mechanochemical cycle detached from actin, sustained force generation emerges only when many motors function collectively within an ensemble (1–4, 7). In this collective setting, force generation is not simply the sum of individual motor properties but instead represents an emergent behavior arising from mechanical coupling, load sharing, and coordination among many interacting motors (2–4, 7–17). However, it is still unclear how mechanically coupled myosin ensembles collectively regulate force generation in response to changes in their mechanical state.

Mechanical feedback plays a key role in ensemble function. External load alters actomyosin kinetics, attachment lifetime, and thus cooperative force sharing among neighboring motors (16–23) (16–21). However, the mechanical environment experienced by each motor is also determined by the ensemble itself. Motor density, filament compliance, crosslinking, and substrate mechanics influence how strain is transmitted through the actomyosin system, thereby shaping collective force generation.(10–12, 14, 15, 20, 22–25). These observations suggest that myosin ensembles continuously adapt to changes in their mechanical environment through feedback between individual motor activity and ensemble mechanics.

Our previous studies using reconstituted skeletal myosin ensembles demonstrated that force generation depends on the local motor environment (13,14,16). Within compliant actin filament bundles, reducing motor density increased coordinated force generation within optical trapping assays, whereas increasing motor density beyond an intermediate level reduced force output despite the presence of additional motors (15, 17). These findings suggested that maximal force generation is achieved through an optimal balance between motor concentration and mechanical coordination, and more motors are not necessarily better in terms of productive force generation (7, 9, 15, 17, 26–30). We further proposed that the actin filament functions as a mechanically responsive element that couples individual motors through emergent mechanical feedback (9–11,14,16,21). In this framework, changes in motor binding alter the mechanical state of the actin filament, while changes in filament mechanics feedback to regulate the attachment and force-producing behavior of neighboring motors. Such a mechanism provides a potential means by which mechanically coupled ensembles coordinate force generation without direct communication between individual motors.

Whether these ensemble-level principles extend to cardiac myosin remains unknown. Although skeletal and cardiac myosin share a common actomyosin mechanochemical cycle, cardiac myosin exhibits slower kinetics, a higher duty ratio, and distinct regulatory mechanisms that govern the availability of force-generating heads (26, 31–42). These differences raise the possibility that cardiac myosin ensembles coordinate force generation differently from skeletal ensembles. More fundamentally, it remains unknown whether changes in effective motor occupancy regulate emergent force generation through the same mechanical feedback mechanisms observed in skeletal myosin.

Full-length cardiac myosin contains two motor heads connected to a coiled-coil tail, allowing interactions between the heads and between the heads and tail that contribute to autoinhibited conformations associated with the interacting-heads motif and the super-relaxed (SRX) state (30, 53, 57, 58). These structural features may also influence the mechanical environment of an ensemble by introducing additional sites of interaction and potential sources of steric interference, drag, and mechanical constraint (17, 43). By contrast, subfragment 1 (S1) retains a single motor head and its associated light chains but lacks the second head and tail. Skeletal muscle myosin S1 has been shown to induce actin filament bundling (44), and cardiac myosin S1 has likewise been shown to form actin bundles (45). Miyanishi et al. identified an additional actin-binding interaction associated with the essential light chain (ELC), beyond the canonical actin-binding interface of the motor domain. Subsequent studies have implicated the ELC N-terminal extension in actin interactions and in the regulation of actomyosin force generation and kinetics (46–51). In particular, Miller et al. proposed that interaction between residues 5–14 of the cardiac ELC N-terminal extension and the C-terminal region of actin inhibits reversal of the myosin power stroke, potentially increasing force maintenance (48). These findings suggest that cardiac S1 can organize actin filaments despite lacking the second head and coiled-coil tail of full-length myosin and that ELC-mediated interactions may influence both filament organization and the mechanical response of actomyosin assemblies. This is further demonstrated as removal of the N-terminus of the ELC produced approximately 30% deficit in myosin crossbridge content in the myocardium of mice (46) and mutations in the ELC have been implicated in HCM (52, 53). However, the extent to which ELC-mediated interactions contribute to force transmission within mechanically coupled actomyosin assemblies remains unclear (53). Comparing full-length and S1 myosin provides a means of investigating how motor loading and myosin architecture influence mechanical feedback and force generation within actin-associated assemblies.

In addition, small-molecule cardiac myosin modulators provide an opportunity to investigate how changes in motor occupancy regulate emergent collective force generation (37, 54–61). Omecamtiv mecarbil (OM) prolongs the lifetime of strongly bound actomyosin interactions, enhances the duty ratio of cardiac myosin, and has been used in heart failure models to increase contractility (54, 58, 62–65). In contrast, mavacamten (MAVA) stabilizes the SRX state, reducing the number of available force-generating myosin heads, and is used clinically to reduce hypercontractility in hypertrophic cardiomyopathy (32, 37–39, 55, 60, 61, 66–69). MAVA has also been found to stabilize the ADP-P_i_ binding state which stalls the motor domain in a primed state and reduces motor dynamics (70). Although these compounds act through distinct molecular mechanisms and produce different physiological outcomes, both perturb factors expected to influence effective motor engagement within an ensemble, including actin-attachment lifetime and the availability of force-generating motors (56, 59, 62, 63, 66). In addition, structural studies have shown that OM and MAVA bind within the same allosteric pocket in the motor domain, despite producing distinct functional effects on cardiac myosin (71) (Houdusse et al., 2024). This shared binding site provides a direct connection between the pharmacological perturbations and the comparison between full-length myosin with S1. The consequences of perturbing motor kinetics or availability may depend on how motor activity is coupled to the surrounding actin network and to other structural elements within the assembly.

To better understand the interplay between motor loading, pharmacological modulation, and myosin architecture in collective force generation, we investigated reconstituted cardiac myosin assemblies containing either full-length (FL) myosin or S1. We varied myosin loading and applied OM or MAVA to investigate how changes in motor availability and kinetics influence collective force generation across different ensemble structures. Our results show that reducing myosin loading increases maximum ensemble force under drug free conditions and that pharmacological responses depend on loading state, with significant loading-dependent effects observed in S1 ensembles. These findings support an occupancy-coordination framework in which collective force generation reflects interactions among motor loading, motor kinetics, and ensemble organization rather than motor number alone.

## Methods

### Actin Polymerization and Protein Reconstitution

Actin filament polymerization was performed as described previously (15, 17, 30, 37, 38, 58, 59, 62, 72). Non-labeled bovine cardiac muscle actin (Cytoskeleton) was reconstituted by adding 100 µL of reverse osmosis (RO) water to 1 mg of lyophilized actin and gently mixing by pipetting. The solution was aliquoted and stored at −80 °C at a final actin concentration of 10 mg/mL. Non-labeled actin was polymerized into filaments by first mixing 5 µL of 10 mg/mL with 50 µL General Actin Buffer (GAB: 5 mM Tris–HCl, 0.2 mM CaCl₂, 0.5 mM DTT, 0.2 mM ATP). The mixture was kept in ice and incubated for 1 h. Actin was then polymerized by adding 5.5 µL Actin Polymerization Buffer (APB: 50 mM Tris-HCl, 500 mM KCl, 2 mM MgCl_2_, 2 mM CaCl_2_, 2 mM DTT, 5 mM ATP), gently mixing by pipetting, and incubating for 20 min on ice. Phalloidin was used to stabilize and stain the polymerized actin filaments by adding 5 µL of rhodamine-labeled phalloidin (Cytoskeleton), wrapping the vial in aluminum foil to block room light, and incubating on ice for 1 h. The labeled actin was stored at 4°C and used within one week to prepare actin–myosin bundles (17, 30, 37, 38, 58, 59, 62, 72).

Biotinylated skeletal muscle actin (from Cytoskeleton) was reconstituted by adding 20 µL RO water to 20 µg of lyophilized actin and gently mixing by pipetting. The solution was aliquoted and stored at −80 °C at a final concentration of 1 mg/mL. To form a 10:1 mixture of unlabeled actin to biotinylated actin, 5 µL of 10 mg/mL unlabeled actin were combined with 5 µL of 1 mg/mL biotinylated actin. The solution was mixed with 100 µL GAB and incubated for 1 h on ice. Biotinylated actin filaments were polymerized by adding 11 µL APB, gently mixing, and incubating for 20 min on ice. Filaments were stabilized and fluorescently labeled by adding 5 µL Alexa Fluor 488–phalloidin (from ThermoFisher), protecting from light, and incubating on ice for 1 h. Labeled filaments were stored at 4°C and used within one week (17, 30, 37, 38, 58, 59, 62, 72). Full-length bovine cardiac myosin II (from Cytoskeleton) was reconstituted to 10 mg/mL by adding 100 µL RO water containing 1 mM DTT, gently mixing, aliquoting, and storing at −80 °C. Working dilutions were prepared fresh for each experiment as specified below. Cardiac myosin S1 (from Cytoskeleton) was reconstituted by briefly centrifuging the lyophilized protein down and dissolved in Milli-Q water (75 µL for 250 µg, 300 µL for 1 mg, or 1.5 mL for 5 mg), yielding a 3.3 mg/mL stock aliquoted and stored at −80 °C (17, 30, 37, 38, 58, 59, 62, 72).

### Actomyosin Bundle Assay Preparation

Optical trapping assay preparation followed the previously published actomyosin bundle protocol, adapted here for cardiac myosin (15, 17). Streptavidin-coated polystyrene beads (Spherotech, 1 µm) were cleaned by diluting 20 µL beads into 80 µL RO water and washing four times by centrifugation at 10,000 rpm. After each wash, beads were resuspended in 100 µL RO water (or GAB for a final wash). Beads were sonicated for 2 min at 40% amplitude and stored on a rotator at 4°C until use (15, 17, 30).

Coverslips were cleaned using an ethanolic potassium hydroxide (KOH) treatment before surface functionalization as described previously (15, 17). Briefly, 100 g of KOH was dissolved in 300 mL of ethanol. Coverslips were placed in Teflon racks, and the KOH–ethanol solution, a separate ethanol bath, and two deionized-water baths were degassed for 5 min using a bath sonicator. Coverslips were immersed in the KOH–ethanol solution and sonicated for 5 min, followed by successive rinsing in ethanol and deionized water until the liquids flowed uniformly across the surfaces without beading. The coverslips were then sonicated in fresh deionized water for an additional 5 min, rinsed thoroughly with deionized water and ethanol, and oven-dried for at least 15 min before poly-L-lysine coating. Etched coverslips were coated by soaking in poly-L-lysine solution (PLL: 30 mL 100% ethanol with 200 µL of 0.1% w/v poly-L-lysine; Sigma-Aldrich) for 15 min to facilitate filament binding. Coverslips were removed with tweezers, holding only the edges, and dried with a filtered airline until no ethanol remained and the surface was residue-free. Flow channels (10–15 µL) were assembled using a PLL-coated coverslip, a microscope slide, and double-sided tape. Two parallel strips of tape were placed on the slide (3–4 mm spacing), and the coverslip was placed perpendicular to the long axis of the microscope slide to form a sealed channel. Pressure was applied to seal the channel using a microtube until the tape became transparent (15, 17, 30).

Actomyosin bundles for optical trapping experiments were formed as described in Al Azzam et al. (15, 17) to probe the mechanics of full-length and S1 cardiac myosin II motor ensembles interacting with actin filaments (15, 17, 30). This assay configuration, as shown in Figure 1, incorporates actin filament structural hierarchy and compliance as an environment for interrogating cardiac myosin ensemble behavior, as opposed to previous single-molecule or small ensemble experiments where motors were immobilized to a rigid surface, such as a bead or coverslip (2–4, 10, 11, 13, 16). To prepare the bundle assay, rhodamine-labeled actin and biotinylated Alexa Fluor 488 actin filament solutions were diluted 600X in APB. To ensure efficient fluorescent labeling, 5 µL of the corresponding labeled phalloidin was added to each tube, and samples were incubated on ice in the dark for 15 min before use. 15 µL of diluted rhodamine-actin was introduced into the PLL-coated flow cell and incubated for 10 min in a humidified chamber. The flow cell was then blocked with 20 µL of 1 mg/mL casein (from Bio-Rad) in APB for 5 min to minimize nonspecific binding. In a separate tube, 15 µL of diluted biotinylated actin was mixed with an oxygen-scavenging system consisting of 1 µL β-D-glucose (500 mg/mL), 1 µL glucose oxidase (25 mg/mL), and 1 µL catalase (500 U/mL), together with 1 µL ATP (100 mM) and 1 µL of cleaned streptavidin beads. This mixture was kept at 4°C in the dark while surface-bound rhodamine-labeled actin filaments were incubated. Immediately before loading into the flow cell, the biotinylated Alexa Fluor 488-labeled actin–bead mixture was mixed with cardiac myosin II at the desired concentration by gentle pipetting. Myosin was added to the biotinylated actin–bead mixture immediately before introduction into the flow cell rather than being preincubated with this mixture. The resulting solution was promptly introduced into the flow channel and incubated for 20 min to allow the formation of mechanically coupled actomyosin assemblies containing surface-associated rhodamine-labeled actin filaments, biotinylated Alexa Fluor 488-labeled actin filaments, and cardiac myosin. These assays are prepared in APB with high ionic strength to prevent thick filament assembly and measure the effect of non-aggregated motors within the bundles (15, 17). The reported cardiac myosin concentrations (0.2 and 0.04 µM) refer to the solution concentrations introduced during actomyosin bundle assembly and were used as experimental proxies for relative myosin loading. The absolute number of myosin molecules incorporated into each individual bundle, and the fraction of incorporated motors that were mechanically active, were not directly measured. Accordingly, comparisons between the two concentrations are interpreted as differences in ensemble loading rather than as defined numbers of active motors.

**Figure 1.**
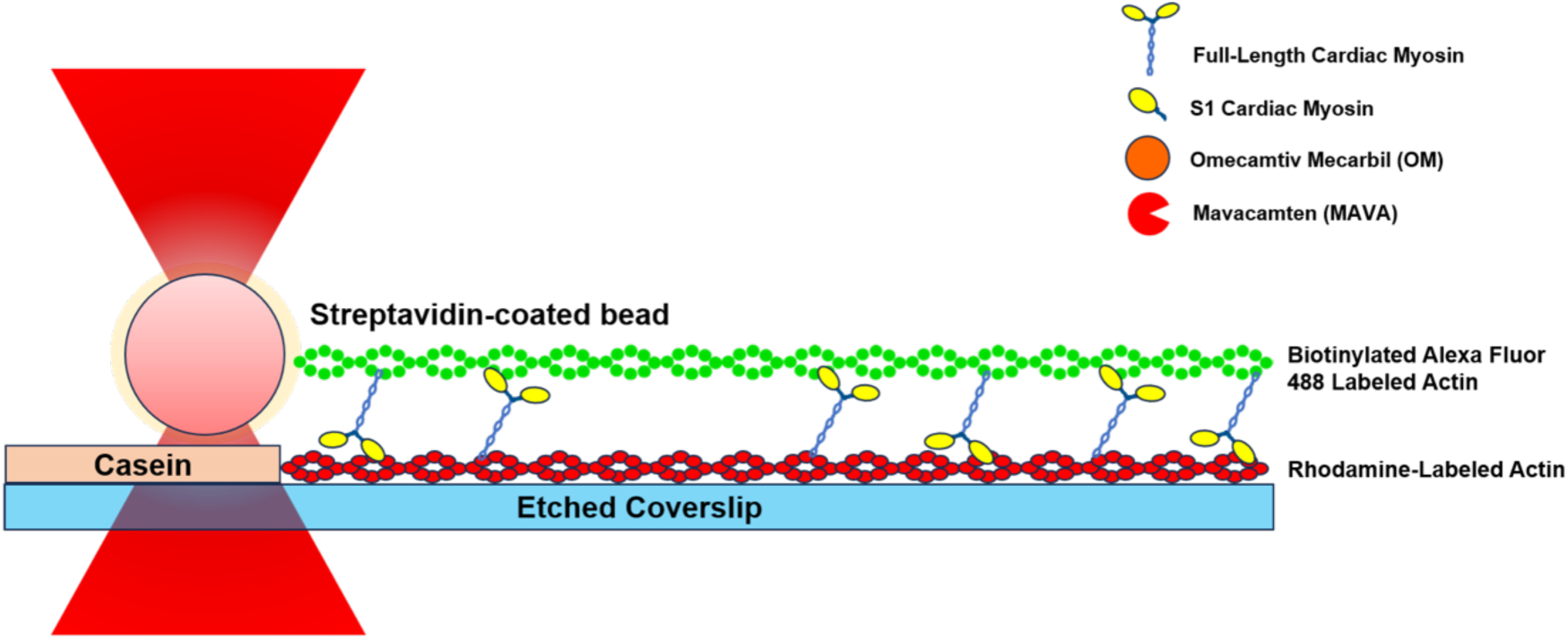
Schematic of the optical tweezers assay used to measure force generation by cardiac myosin ensembles. Rhodamine-labeled actin filaments were adsorbed onto a poly-L-lysine-coated etched coverslip, and casein was used to block nonspecific interactions. Myosin constructs, biotinylated Alexa Fluor 488-labeled actin filaments, and streptavidin-coated beads were then added together, allowing actomyosin assemblies to self-assemble on the coverslip surface. Force generating assemblies formed through interactions between myosin and two differently labeled actin filaments, with the bead-associated filament providing a handle for optical trapping measurements. Force generation was observed with both FL myosin and S1, which contains the myosin motor domain but lacks the tail. The assay was performed under elevated ionic strength conditions to limit myosin thick filament formation and investigate how myosin motor ensembles self-organize to support force generation. Following assembly formation, omecamtiv mecarbil (OM) or mavacamten (MAVA) was introduced to examine how pharmacological modulation of motor activity influences force generation and ensemble mechanics.

Following the 20 min incubation, for control (drug-free) slides, the channel was washed with a solution containing 30 µL APB and 1 µL streptavidin beads, with no drug, and then sealed with nail polish to prevent evaporation during experiments. For drug-treated slides, the final wash was prepared by mixing 30 µL APB, 1 µL streptavidin beads, and the appropriate amount of OM or MAVA (purchased from Selleck Chemicals (Houston, TX, USA)). OM and MAVA powders were dissolved in DMSO to prepare 0.4, 0.8, and 1.6 mg/mL stocks, which were aliquoted and stored at −80 °C. Working solutions were prepared fresh by dilution into APB to the desired final in-channel concentration. This drug-containing wash was introduced immediately after the incubation step to remove unbound material and deliver the drug compound. The channel was then sealed with nail polish to prevent evaporation during the experiment. APB served as the base buffer for the final acquisition wash. Following addition of 1 µL bead suspension to 30 µL APB, the acquisition solution contained approximately 4.88 mM total ATP, 1.84 mM total MgCl₂, and 1.84 mM total CaCl₂. Mg–ATP complex formation and free Mg²⁺ were estimated using PyChelator v1.2.0 with NIST stability constants at 21 °C, nominal pH 7.5, and an ionic equivalent of 0.50 M, yielding approximately 1.64 mM Mg–ATP and 0.20 mM free Mg²⁺. No additional MgCl₂ was added during force acquisition (73).

### Optical Trapping Measurements and Fluorescence Imaging

Flow cells containing the actomyosin bundles were imaged and measured using a JPK/Bruker Nano Tracker 2 optical-trapping instrument equipped with DIC and epifluorescence imaging (**Figure 1**). Epifluorescence imaging of actin filaments was achieved by excitation with a Photofluor LM-75 lamp in conjunction with 488 and 532 nm filter cubes. Actomyosin assemblies were identified through fluorescence imaging confirmation of colocalization between surface-bound rhodamine-labeled actin filaments and bead-associated biotinylated Alexa Fluor 488-labeled actin filaments, as well as their alignment along the same parallel axis, following the previously established assay protocol (15, 17). The polarity of the actin filaments within individual assemblies were not known, and the measured bead displacement was not assigned to a specific direction relative to actin filament polarity. Optical trap calibration was performed at the beginning of each experiment using a freely suspended 1µm streptavidin-coated bead introduced during the final bead-containing wash. Calibration was conducted using the power-spectrum routine implemented in the JPK/Bruker NanoTracker 2 software at a sampling frequency of 3000 Hz. The average trap stiffness was 0.03 pN/nm across compiled calibration records from each experiment. After bead calibration, a bead bound to the biotinylated 488-labeled actin filament within a confirmed bundle was then optically trapped. The beginning of each analyzed trace (t = 0 s) corresponded to initiation of force-data acquisition in the JPK NanoTracker software after a bead associated with a confirmed actomyosin bundle had been captured in the optical trap. Force recordings were evaluated over a standardized 400-s observation window. To ensure measurements were not associated with drift, the position of a streptavidin bead non-covalently attached to an etched coverslip coated in poly-l-lysine was recorded over 400 s, matching the duration of the force measurements. The resulting traces exhibited high noise, and no significant force generation was detected (<0.03 pN). In contrast, tighter noise observed in the actomyosin bundle traces is consistent with reduced Brownian motion through engagement of the bead with force generating actomyosin assemblies (74, 75). Custom MATLAB code was used to visualize traces and perform position and force analyses. Change in position per time was used to calculate force and mean force for each condition (motor concentration, myosin construct, and the presence or absence of drug). The force trajectories obtained in this reconstituted assay represent the time-dependent mechanical output of the assembled actomyosin system and may reflect contributions from motor-generated force together with ensemble-scale structural rearrangement during force development. Because several traces evolved over hundreds of seconds, the measured temporal behavior is interpreted as the dynamics of the reconstituted ensemble assay and not as a direct representation of physiological cardiac contraction kinetics.

### Statistical Analysis

Statistical analyses were performed in MATLAB (MathWorks, Natick, MA, USA) using the Anova function. Force recordings were analyzed over a standardized observation window of 400 s, beginning at the initiation of force-data acquisition. In a few cases, free floating beads would approach the trapped bead during an experiment, and to prevent the bead from falling into the laser trap and disrupting the measurement, the measurement would be stopped before reaching 400 s. For each independent actomyosin assembly, the primary quantitative endpoint was defined as the maximum bead-displacement-derived force attained within this interval. This metric represents the largest measured force within the specified observation window and does not necessarily correspond to a mechanical plateau, an isometric force, or the stall force of an individual motor or motor ensemble. Each independent actomyosin ensemble contributed one maximum-force measurement to the statistical analysis. Each independent actomyosin ensemble contributed one maximum-force measurement to the statistical analysis. Baseline experiments comparing full-length (FL) and S1 cardiac myosin were analyzed using two-way analysis of variance (ANOVA) with myosin construct (FL or S1) and myosin concentration (0.2 or 0.04 µM) as fixed factors. Experiments evaluating omecamtiv mecarbil (OM) or mavacamten (MAVA) were analyzed separately for each construct using two-way ANOVA with myosin concentration (0.2 or 0.04 µM) and drug concentration (no drug, 0.4 mg/mL, or 0.8 mg/mL) as fixed factors. Interaction terms between the two factors were included in all models to determine whether drug responses depended on myosin concentration. Type III sums of squares were used because sample sizes differed among experimental conditions. Representative force traces are presented to illustrate qualitative differences in the temporal evolution, persistence, and apparent coordination of force generation across independent actomyosin ensembles. These dynamic trace features were not used as quantitative statistical endpoints in the present analysis. ANOVA therefore evaluated differences in the maximum force attained within the standardized 400-s observation window. Results are reported as F-statistics with corresponding degrees of freedom and p-values. Statistical significance was defined as p < 0.05.

## Results

### Collective force generation is enhanced at reduced myosin concentration in the absence of drug

To establish how motor loading influences collective force generation, the maximum force generated by full-length (FL) and S1 cardiac myosin ensembles was quantified over a standardized 400-s observation window at two myosin concentrations in the absence of pharmacological modulators. Myosin concentrations of 0.2 and 0.04 µM refer to the solution concentrations introduced into the flow cell during actomyosin bundle assembly. These measurements quantify the maximum force transmitted to the trapped bead during the defined observation period and are not interpreted as steady-state isometric or stall forces. Because individual trajectories did not necessarily reach a plateau within 400 s, the reported maximum-force values describe the observed mechanical responses within this common measurement window rather than an equilibrium force-generating capacity.

Reduced myosin loading was associated with greater maximum force generation in both FL and S1 ensembles (**Figure 2**). Mean maximum force increased from 0.43 ± 0.31 pN at 0.2 µM to 1.51 ± 2.06 pN at 0.04 µM for FL ensembles and from 0.32 ± 0.14 pN to 1.02 ± 0.60 pN for S1 ensembles. Two-way ANOVA identified a significant main effect of myosin concentration on maximum force (F₁,₂₅ = 4.28, p = 0.049), whereas neither myosin construct (F₁,₂₅ = 0.48, p = 0.495) nor the construct × myosin concentration interaction (F₁,₂₅ = 0.19, p = 0.664) was significant. Thus, reduced myosin loading was associated with greater maximum force generation under no-drug conditions in both FL and S1 ensembles. Substantial variability was observed between individual force measurements, particularly in the reduced-concentration full-length myosin condition. However, all measurements were retained in the primary analysis because no independent evidence of experimental failure was identified.

**Figure 2.**
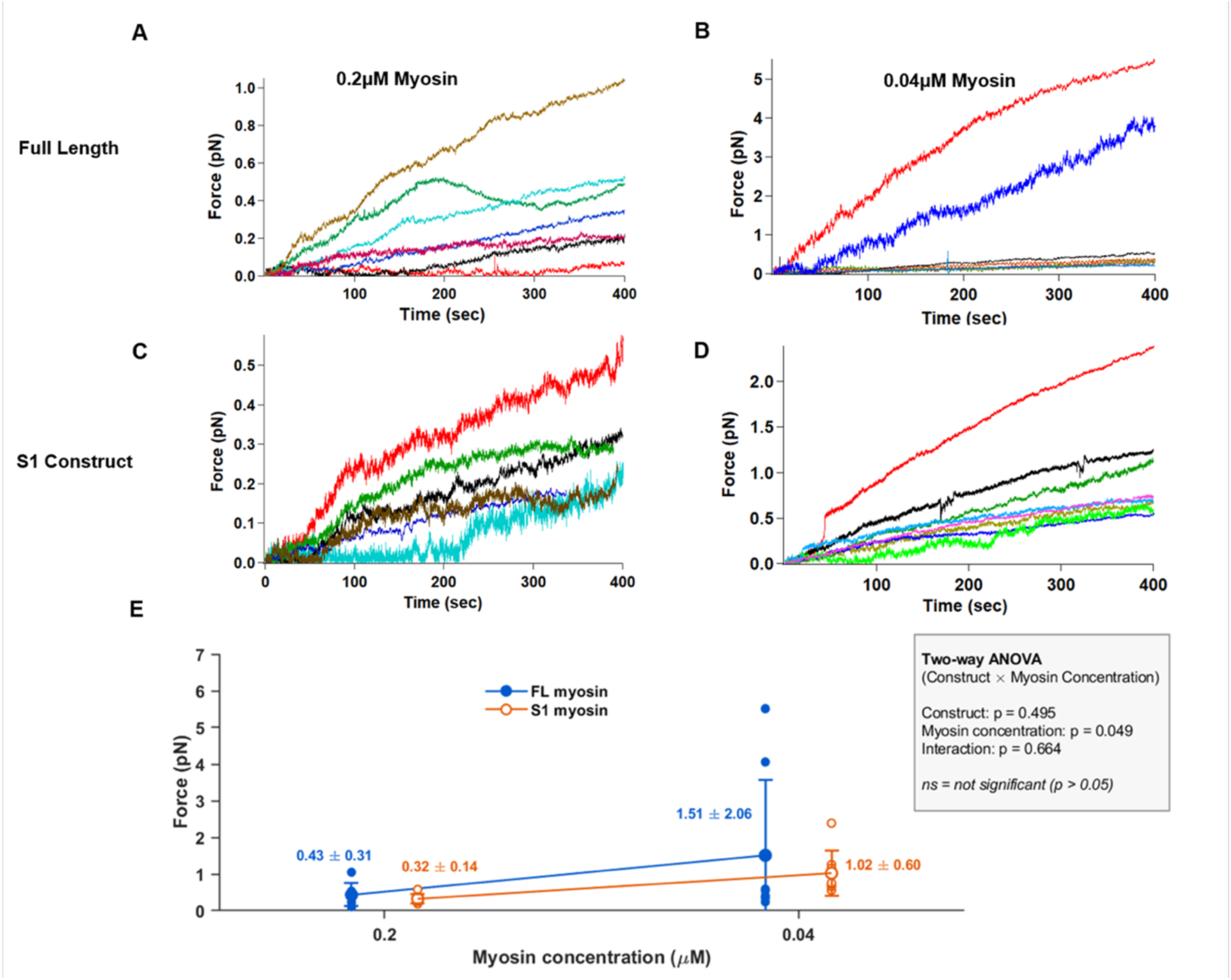
Myosin loading concentration influences endpoint force generation in cardiac myosin ensembles in the absence of pharmacological modulation. **(A,B)** Representative force– time traces generated by full-length (FL) cardiac myosin ensembles assembled using 0.2 µM **(A)** or 0.04 µM **(B)** myosin solution concentration in the absence of drug. (C, D) Representative force– time traces generated by S1 cardiac myosin ensembles assembled using 0.2 µM **(C)** or 0.04 µM **(D)** myosin solution concentration in the absence of drug. The indicated concentrations refer to the myosin solution concentrations introduced into the flow cell for actomyosin bundle assembly. Reducing myosin concentration from 0.2 µM to 0.04 µM produced smoother force trajectories and increased the occurrence of higher-force events in both FL and S1 ensembles. **(E)** Summary of maximum force measurements within the 400-s observation window for FL and S1 ensembles under no-drug conditions. Smaller circles represent individual trace maximum-force measurements, and larger circles with error bars indicate mean ± SD. Mean maximum forces were 0.43 ± 0.31 pN for FL 0.2 µM (n = 7), 1.51 ± 2.06 pN for FL 0.04 µM (n = 8), 0.32 ± 0.14 pN for S1 0.2 µM (n = 6), and 1.02 ± 0.60 pN for S1 0.04 µM (n = 8). Two-way ANOVA identified a significant main effect of myosin concentration [F (1,25) = 4.28, p = 0.049], but no significant effect of construct [F (1,25) = 0.48, p = 0.495] or construct × concentration interaction [F (1,25) = 0.19, p = 0.664]. FL, full-length; S1, myosin subfragment 1.

Representative force–time traces also revealed qualitative differences in force fluctuations between loading conditions (Figure 2). At reduced myosin concentration, trajectories generally appeared smoother, with less pronounced short-timescale fluctuations, particularly in S1 ensembles. In contrast, trajectories at the higher myosin concentration exhibited more frequent and pronounced fluctuations. These features were not quantitatively analyzed and are therefore presented as qualitative characteristics of the representative trajectories. Together, the maximum-force measurements and representative traces indicate that changing myosin loading influenced the magnitude of collective force generation and was accompanied by qualitative differences in temporal behavior.

### Omecamtiv mecarbil alters force generation in a myosin concentration-dependent manner

To determine how increasing actin-attachment lifetime influences collective force generation, FL and S1 myosin ensembles were exposed to increasing concentrations of omecamtiv mecarbil (OM). Mean force responses and representative force trajectories varied across both myosin loading and OM concentration, although statistically significant loading-dependent effects were observed only in S1 ensembles.

For FL ensembles, mean maximum force measured within a bundle’s force trace differed across OM concentrations at the two myosin loading conditions (Figure 3). At 0.2 µM FL myosin, mean maximum force increased from 0.43 ± 0.31 pN without drug to 1.03 ± 0.89 pN at 0.4 mg/mL OM before decreasing to 0.40 ± 0.30 pN at 0.8 mg/mL OM. In contrast, at 0.04 µM FL myosin, mean maximum force decreased from 1.51 ± 2.06 pN without drug to 0.40 ± 0.18 pN at 0.4 mg/mL OM, followed by an increase to 0.78 ± 0.44 pN at 0.8 mg/mL OM. However, two-way ANOVA detected no significant main effect of myosin concentration (F₁,₃₉ = 0.86, p = 0.359), OM concentration (F₂,₃₉ = 0.62, p = 0.541), or myosin concentration × OM concentration interaction (F₂,₃₉ = 2.49, p = 0.096). Thus, although the FL group means displayed different response patterns across loading conditions, the maximum-force analysis did not establish a statistically significant loading-dependent effect of OM.

**Figure 3.**
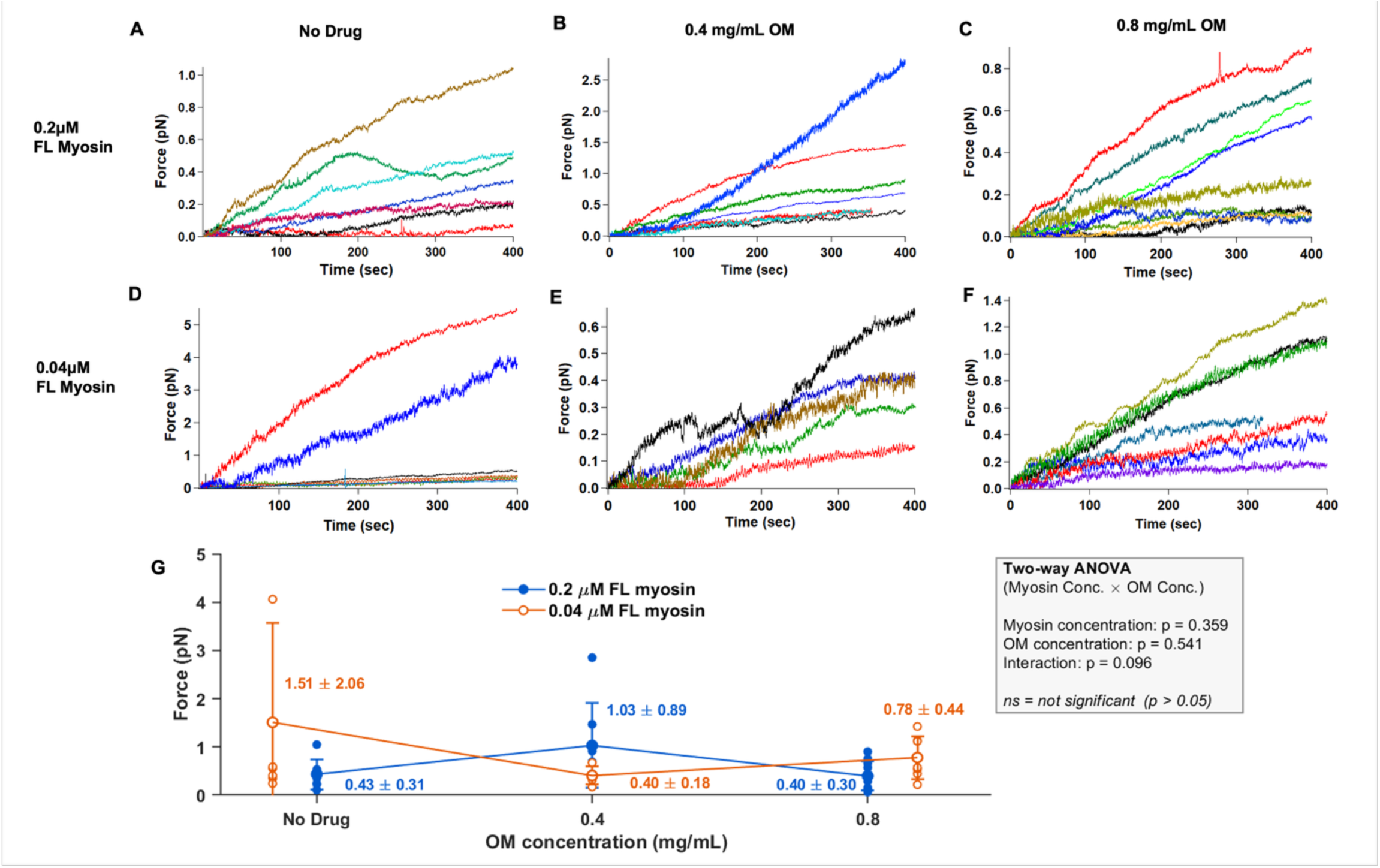
Omecamtiv mecarbil alters force-generation patterns in full-length cardiac myosin ensembles across myosin loading conditions. >**(A–C)** Representative force–time traces generated by full-length (FL) cardiac myosin ensembles assembled using 0.2 µM myosin in the absence of drug (A), with 0.4 mg/mL OM (B), or with 0.8 mg/mL OM (C). **(D–F)** Representative force–time traces generated by FL cardiac myosin ensembles assembled using 0.04 µM myosin in the absence of drug **(D)**, with 0.4 mg/mL OM **(E)**, or with 0.8 mg/mL OM **(F)**. The indicated myosin concentrations refer to the solution concentrations introduced into the flow cell for actomyosin bundle assembly. **(G)** Summary of maximum force measured within the standardized 400-s observation window. Smaller circles represent individual trace measurements, and larger circles with error bars indicate mean ± SD. All individual force measurements were retained in the primary analysis, including the two higher-force observations in the 0.04 µM FL no-drug condition. These measurements contribute to the substantial variability observed in this condition and influence the reported group mean. At 0.2 µM myosin, mean maximum force was 0.43 ± 0.31 pN without drug (n = 7), 1.03 ± 0.89 pN with 0.4 mg/mL OM (n = 7), and 0.40 ± 0.30 pN with 0.8 mg/mL OM (n = 11). At 0.04 µM myosin, mean maximum force was 1.51 ± 2.06 pN without drug (n = 8), 0.40 ± 0.18 pN with 0.4 mg/mL OM (n = 5), and 0.78 ± 0.44 pN with 0.8 mg/mL OM (n = 7). Two-way ANOVA identified no significant main effect of myosin concentration [F (1,39) = 0.86, p = 0.359], no significant main effect of OM concentration [F (2,39) = 0.62, p = 0.541], and no significant myosin concentration × OM concentration interaction [F (2,39) = 2.49, p = 0.096]. OM, omecamtiv mecarbil; FL, full-length.

Representative FL force trajectories also showed qualitative differences across OM conditions (**Figure 3**). At 0.2 µM FL myosin, the OM-treated trajectories remained predominantly ramp-like, with differences in the spread and magnitude of force development across drug concentrations. At 0.04 µM FL myosin, OM treatment was accompanied by more pronounced short-timescale irregularity, particularly at 0.4 mg/mL, relative to the comparatively smoother no-drug trajectories. These features were not quantitatively analyzed and are therefore reported descriptively; however, the representative trajectories suggest that OM treatment was associated with changes in force dynamics not captured by maximum force alone.

S1 ensembles exhibited a statistically significant loading-dependent response to OM (Figure 4). At 0.2 µM S1 myosin, mean maximum force increased from 0.32 ± 0.14 pN without drug to 1.10 ± 0.79 pN at 0.4 mg/mL OM and 1.40 ± 1.12 pN at 0.8 mg/mL OM. In contrast, at 0.04 µM S1 myosin, maximum force decreased from 1.02 ± 0.60 pN without drug to 0.70 ± 0.36 pN at 0.4 mg/mL OM, followed by an increase to 0.99 ± 0.71 pN at 0.8 mg/mL OM. Two-way ANOVA revealed a significant myosin concentration × OM concentration interaction (F₂,₄₅ = 3.60, p = 0.035), whereas neither the main effect of myosin concentration (F₁,₄₅ = 0.03, p = 0.866) nor OM concentration (F₂,₄₅ = 1.99, p = 0.149) was significant. These results demonstrate that the effect of OM on S1 maximum force depended on the myosin loading condition.

**Figure 4.**
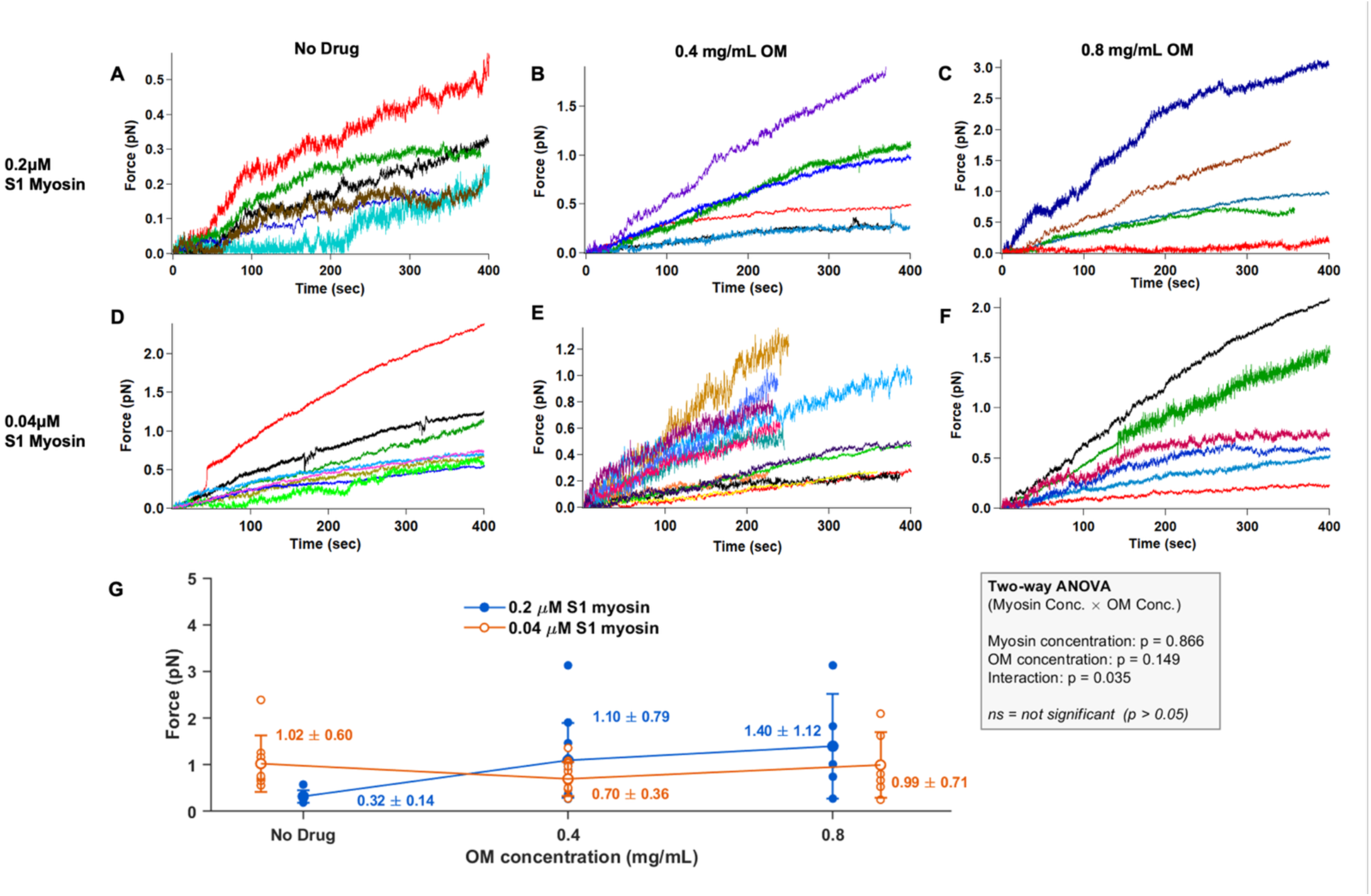
Omecamtiv mecarbil produces myosin-loading-dependent changes in maximum force generation by S1 cardiac myosin ensembles. **(A–C)** Representative force–time traces generated by S1 cardiac myosin ensembles assembled using 0.2 µM myosin in the absence of drug (A), with 0.4 mg/mL OM **(B)**, or with 0.8 mg/mL OM **(C)**. **(D–F)** Representative force–time traces generated by S1 cardiac myosin ensembles assembled using 0.04 µM myosin in the absence of drug **(D)**, with 0.4 mg/mL OM **(E)**, or with 0.8 mg/mL OM **(F)**. The indicated myosin concentrations refer to the solution concentrations introduced into the flow cell for actomyosin bundle assembly. **(G)** Summary of maximum force measured within the standardized 400-s observation window. Smaller circles represent individual trace measurements, and larger circles with error bars indicate mean ± SD. At 0.2 µM myosin, mean maximum force was 0.32 ± 0.14 pN without drug (n = 6), 1.10 ± 0.79 pN with 0.4 mg/mL OM (n = 12), and 1.40 ±1.12 pN with 0.8 mg/mL OM (n = 5). At 0.04 µM myosin, mean maximum force was 1.02 ± 0.60 pN without drug (n = 8), 0.70 ± 0.36 pN with 0.4 mg/mL OM (n = 14), and 0.99 ± 0.71 pN with 0.8 mg/mL OM (n = 6). Two-way ANOVA identified no significant main effect of myosin concentration [F (1,45) = 0.03, p = 0.866] or OM concentration [F (2,45) = 1.99, p = 0.149], but revealed a significant myosin concentration × OM concentration interaction [F (2,45) = 3.60, p = 0.035]. OM, omecamtiv mecarbil; S1, myosin subfragment 1.

Representative S1 force trajectories revealed additional qualitative changes in force fluctuations following OM treatment that differed across myosin loading conditions (**Figure 4**). At the higher S1 concentration, the no-drug condition exhibited pronounced force fluctuations, whereas the addition of OM generally produced smoother trajectories. In contrast, trajectories at the lower S1 concentration appeared comparatively smooth in the absence of drug but exhibited more pronounced fluctuations at 0.4 mg/mL OM. At 0.8 mg/mL OM, these fluctuations appeared reduced relative to the intermediate OM condition in several trajectories. Although these features were not quantitatively analyzed, the representative traces suggest that OM was associated with qualitatively different changes in force dynamics across myosin loading conditions.

### Mavacamten produces biphasic responses at high myosin concentration

To investigate how reducing the availability of force-generating myosin heads influences collective force generation, FL and S1 ensembles were treated with increasing concentrations of mavacamten (MAVA). Mean maximum force that a certain bundle generated during the trace window and the qualitative behavior of representative force trajectories varied across myosin loading and MAVA concentration, with statistically significant loading-dependent effects observed in S1 ensembles.

For FL ensembles, the mean maximum-force response at 0.2 µM myosin was biphasic, increasing at the intermediate MAVA concentration before decreasing at the higher concentration (Figure 5). Mean maximum force increased from 0.43 ± 0.31 pN without drug to 0.86 ± 0.52 pN at 0.4 mg/mL MAVA and then decreased to 0.51 ± 0.41 pN at 0.8 mg/mL MAVA. In contrast, at 0.04 µM FL myosin, mean maximum force was highest in the absence of drug (1.51 ± 2.06 pN) and decreased to 0.82 ± 0.54 pN at 0.4 mg/mL MAVA and 0.69 ± 0.28 pN at 0.8 mg/mL MAVA. Despite these differences in mean responses, two-way ANOVA detected no significant main effect of myosin concentration (F₁,₄₂ = 2.22, p = 0.144), MAVA concentration (F₂,₄₂ = 0.56, p = 0.573), or the myosin concentration × MAVA concentration interaction (F₂,₄₂ = 1.60, p = 0.215). Thus, the biphasic mean response observed at high FL loading and the decrease in mean maximum force observed at low FL loading were not statistically established using maximum force as the endpoint.

**Figure 5.**
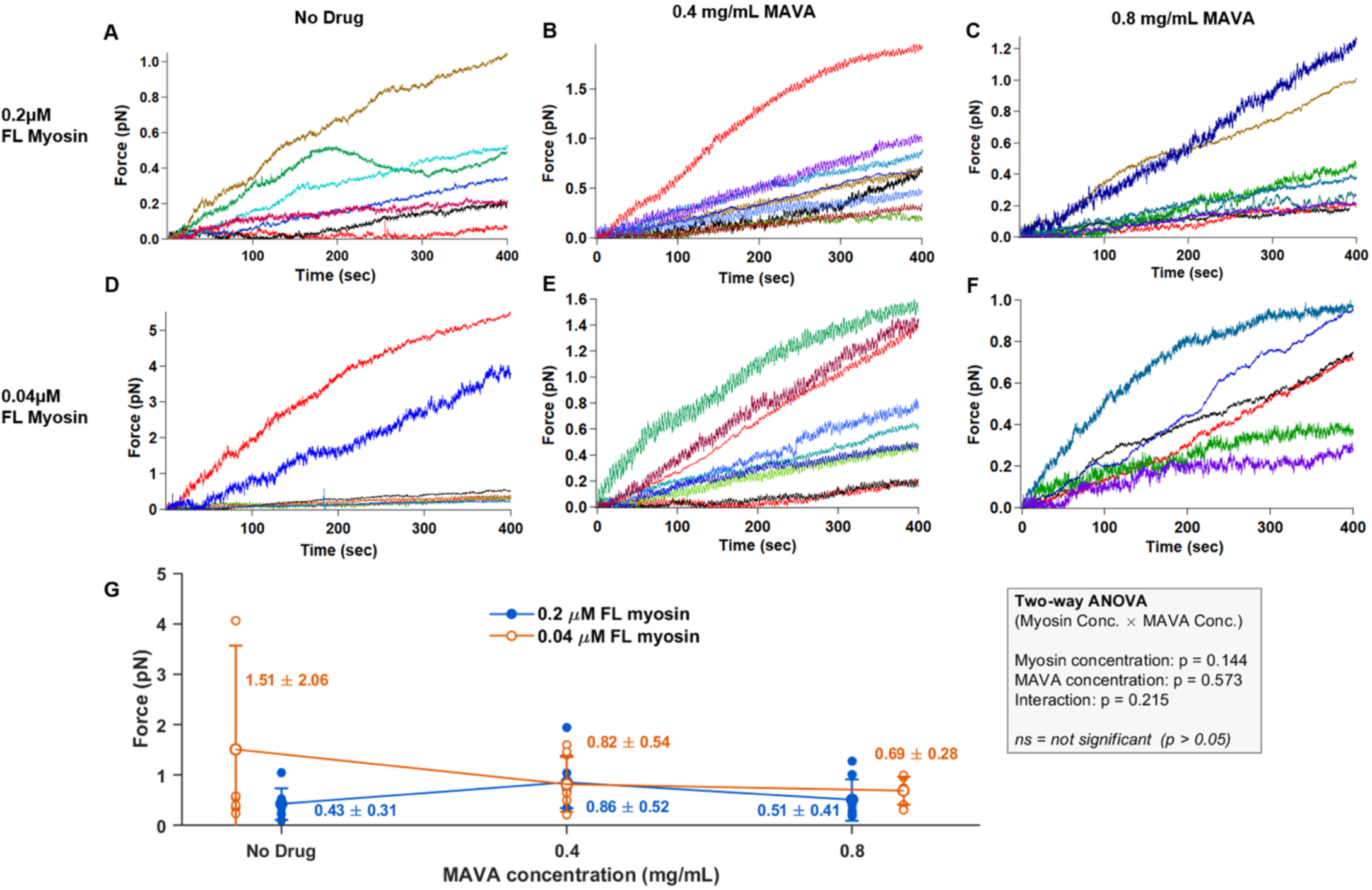
Mavacamten alters force-generation patterns in full-length cardiac myosin ensembles across myosin loading conditions. **(A–C** Representative force–time traces generated by full-length (FL) cardiac myosin ensembles assembled using 0.2 µM myosin in the absence of drug **(A)**, with 0.4 mg/mL MAVA **(B)**, or with 0.8 mg/mL MAVA **(C)**. **(D–F)** Representative force–time traces generated by FL cardiac myosin ensembles assembled using 0.04 µM myosin in the absence of drug **(D)**, with 0.4 mg/mL MAVA **(E)**, or with 0.8 mg/mL MAVA **(F)**. The indicated myosin concentrations refer to the solution concentrations introduced into the flow cell for actomyosin bundle assembly. **(G)** Summary of maximum force measured within the standardized 400-s observation window. Smaller circles represent individual trace measurements, and larger circles with error bars indicate mean ± SD. At 0.2 µM myosin, mean maximum force was 0.43 ± 0.31 pN without drug (n = 7), 0.86 ± 0.52 pN with 0.4 mg/mL MAVA (n = 10), and 0.51 ± 0.41 pN with 0.8 mg/mL MAVA (n = 8). At 0.04 µM myosin, mean maximum force was 1.51 ± 2.06 pN without drug (n = 8), 0.82 ± 0.54 pN with 0.4 mg/mL MAVA (n = 9), and 0.69 ± 0.28 pN with 0.8 mg/mL MAVA (n = 6). Two-way ANOVA identified no significant main effect of myosin concentration [F (1,42) = 2.22, p = 0.144], no significant main effect of MAVA concentration [F (2,42) = 0.56, p = 0.573], and no significant myosin concentration × MAVA concentration interaction [F (2,42) = 1.60, p = 0.215]. MAVA, mavacamten; FL, full-length.

Representative FL trajectories showed qualitative differences across MAVA conditions (**Figure 5**). At 0.2 µM FL myosin, MAVA treatment was accompanied by differences in the spread and time-dependent development of force trajectories, without an obvious uniform increase in short-timescale fluctuations. At 0.04 µM FL myosin, the 0.4 mg/mL MAVA condition displayed more visible short-timescale irregularity than the no-drug condition, whereas the 0.8 mg/mL condition showed a mixture of smoother and more irregular trajectories. These features were not quantified and therefore cannot be compared statistically, but they illustrate qualitative differences in force dynamics across MAVA conditions.

S1 ensembles exhibited a statistically significant loading-dependent response to MAVA (**Figure 6**). At 0.2 µM S1 myosin, mean maximum force increased from 0.32 ± 0.14 pN without drug to 0.66 ± 0.24 pN at 0.4 mg/mL MAVA before decreasing to 0.36 ± 0.33 pN at 0.8 mg/mL MAVA. In contrast, at 0.04 µM S1 myosin, maximum force progressively decreased from 1.02 ± 0.60 pN without drug to 0.77 ± 0.20 pN at 0.4 mg/mL MAVA and 0.41 ± 0.31 pN at 0.8 mg/mL MAVA. Two-way ANOVA revealed a significant main effect of myosin concentration (F₁,₃₃ = 6.11, p = 0.019) and a significant myosin concentration × MAVA concentration interaction (F₂,₃₃ = 3.33, p = 0.048), whereas the main effect of MAVA concentration did not reach statistical significance (F₂,₃₃ = 3.04, p = 0.062). Thus, the effect of MAVA on S1 maximum force depended significantly on the myosin loading condition.

**Figure 6.**
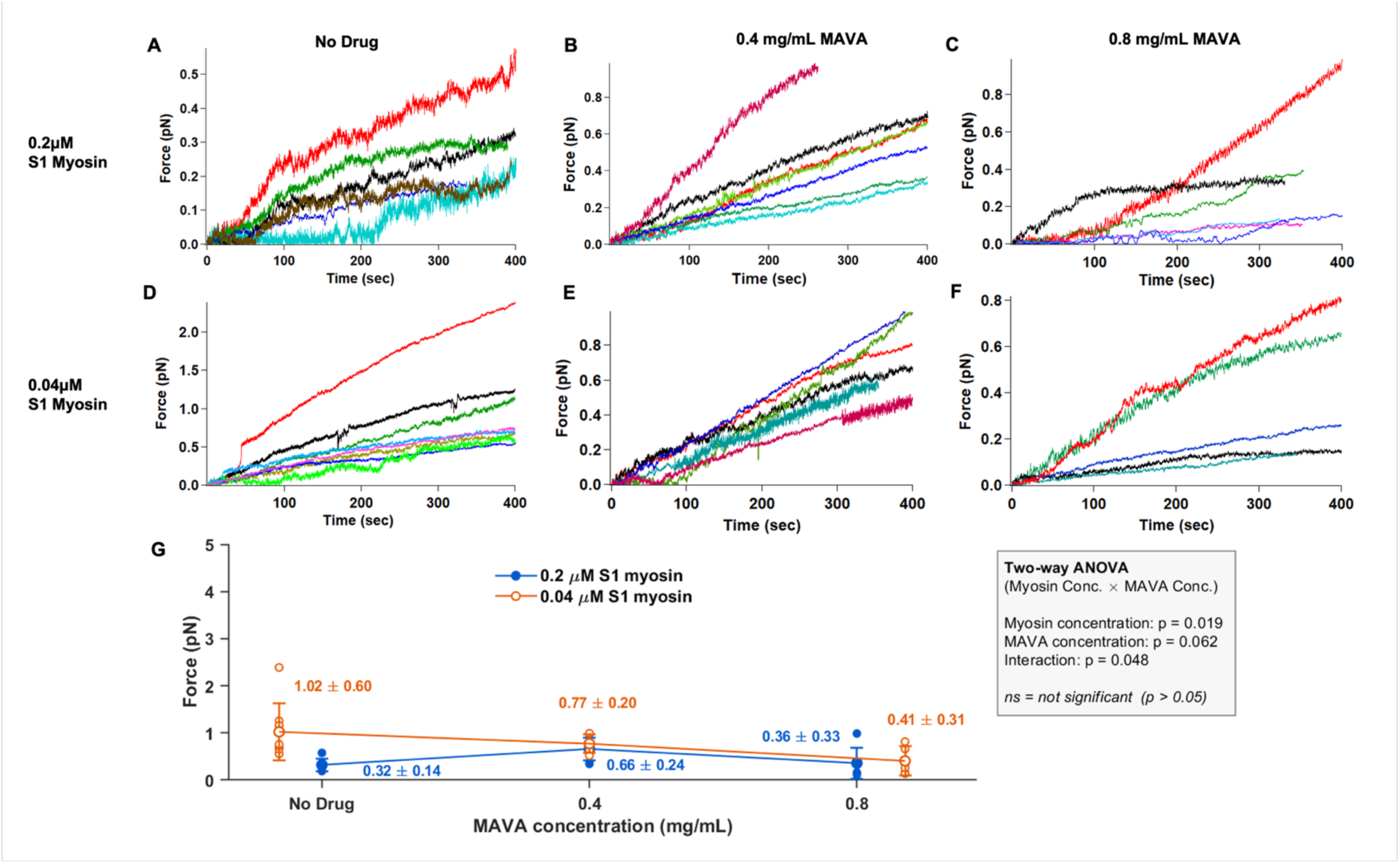
Mavacamten produces myosin-loading-dependent changes in maximum force generation by S1 cardiac myosin ensembles. **(A–C)** Representative force–time traces generated by S1 cardiac myosin ensembles assembled using 0.2 µM myosin in the absence of drug **(A)**, with 0.4 mg/mL MAVA **(B)**, or with 0.8 mg/mL MAVA **(C)**. **(D–F)** Representative force–time traces generated by S1 cardiac myosin ensembles assembled using 0.04 µM myosin in the absence of drug **(D)**, with 0.4 mg/mL MAVA **(E)**, or with 0.8 mg/mL MAVA **(F)**. The indicated myosin concentrations refer to the solution concentrations introduced into the flow cell for actomyosin bundle assembly. **(G)** Summary of maximum force measured within the standardized 400-s observation window. Smaller circles represent individual trace measurements, and larger circles with error bars indicate mean ± SD. At 0.2 µM myosin, mean maximum force was 0.32 ± 0.14 pN without drug (n = 6), 0.66 ± 0.24 pN with 0.4 mg/mL MAVA (n = 8), and 0.36 ± 0.33 pN with 0.8 mg/mL MAVA (n = 6). At 0.04 µM myosin, mean maximum force was 1.02 ± 0.60 pN without drug (n = 8), 0.77 ± 0.20 pN with 0.4 mg/mL MAVA (n = 6), and 0.41 ± 0.31 pN with 0.8 mg/mL MAVA (n = 5). Two-way ANOVA identified a significant main effect of myosin concentration [F (1,33) = 6.11, p = 0.019], no significant main effect of MAVA concentration [F (2,33) = 3.04, p = 0.062], and a significant myosin concentration × MAVA concentration interaction [F (2,33) = 3.33, p = 0.048]. MAVA, mavacamten; S1, myosin subfragment 1.

Representative S1 trajectories also showed qualitative differences in force fluctuations across MAVA and myosin loading conditions (**Figure 6**). At the higher S1 concentration, no-drugtrajectories exhibited pronounced fluctuations that generally decreased following MAVA treatment. At the lower S1 concentration, no-drug trajectories appeared comparatively smooth, whereas MAVA treatment was associated with increased fluctuations in some trajectories. These observations were not quantitatively analyzed and are therefore presented as descriptive features of the representative traces; however, they further illustrate that changes in motor availability were accompanied by qualitatively different force dynamics across initial loading conditions.

Across all experimental conditions, maximum force measured within the standardized 400-s observation window served as the primary quantitative endpoint for statistical comparisons. This analysis identified a significant effect of myosin loading under basal conditions and significant myosin loading × drug interactions for both OM and MAVA in S1 ensembles. In contrast, the mean maximum-force responses observed in FL ensembles did not produce statistically significant loading × drug interactions. In parallel, representative force–time trajectories revealed qualitative differences in force fluctuations, smoothness, persistence, and force development across loading, drug, and construct conditions. Because these dynamic features were not quantitatively analyzed in this study, they cannot be interpreted as statistically established differences. Nevertheless, their recurring variation across experimental conditions suggests that maximum force observed within the trace window captures only one aspect of collective ensemble mechanics and motivates future quantitative analysis of force dynamics beyond this metric alone.

## Discussion

Collective force generation by molecular motor ensembles is often expected to increase with the number of available motors. The results presented indicate that increasing motor loading does not necessarily increase ensemble force output proportionally. Because only two myosin-loading concentrations were examined, the experiments do not directly define the shape of the force–loading relationship. Under no-drug conditions, mean maximum force observed in the trace was greater at reduced myosin loading in both FL and S1 ensembles, and two-way ANOVA identified a significant main effect of myosin concentration across constructs. Thus, increasing the concentration of motors introduced during bundle assembly did not produce a proportional increase in collective force output. Rather, these findings suggest that ensemble performance depends on how motor loading influences collective mechanical organization.

One possible explanation for a non-proportional relationship between motor loading and ensemble force generation is that increasing motor density introduces competing effects. Additional motors may increase the number of potential force-generating interactions with actin but could also increase steric interference, competition for available binding sites, internal mechanical strain, or resistive interactions within the ensemble (10, 13, 17, 19, 76). Under this interpretation, collective force generation may be favored at intermediate levels of effective motor engagement rather than increasing monotonically with motor number. The greater maximum forces observed at reduced myosin loading suggest that higher motor occupancy can become mechanically counterproductive.

Recent theoretical work provides an independent physical basis for such occupancy-dependent collective behavior. Barutel and Fürthauer developed a nonequilibrium thermodynamic framework in which motorized crosslinkers compete for a finite number of available binding sites along cytoskeletal filaments. In their model, motor transport becomes intrinsically density-dependent because the thermodynamic susceptibility governing motor flux depends on both motor density and the remaining available filament space. At sufficiently high density, competition for available sites can oppose motor-driven transport and generate traffic-jam-like accumulation of motors (76). Although this model addresses motor transport and pattern formation rather than cardiac myosin force generation directly, it demonstrates a general principle relevant to the findings where increasing motor density need not produce a proportional increase in productive collective activity. Thus, the greater forces observed at reduced myosin loading are consistent with a broader physical picture in which ensemble performance depends on the relationship between motor occupancy, available interaction space, and collective motor organization.

Within this idea, OM and MAVA can be viewed as shifting ensembles in opposite directions along an effective motor occupancy landscape (**Figure 7**). By prolonging actin attachment, OM increases the fraction of motors that may remain engaged with actin at a given time, thereby increasing motor occupancy (37, 54, 62–65). Conversely, MAVA suppresses cardiac myosin motor activity and, in full-length myosin, stabilizes an autoinhibited SRX state that reduces the availability of force-generating heads (60, 61, 66–68, 77). Under the proposed model, the effects of OM and MAVA reflect not only their distinct molecular mechanisms but also the ensemble’s initial occupancy state and the mechanical constraints under which the motors operate.

**Figure 7.**
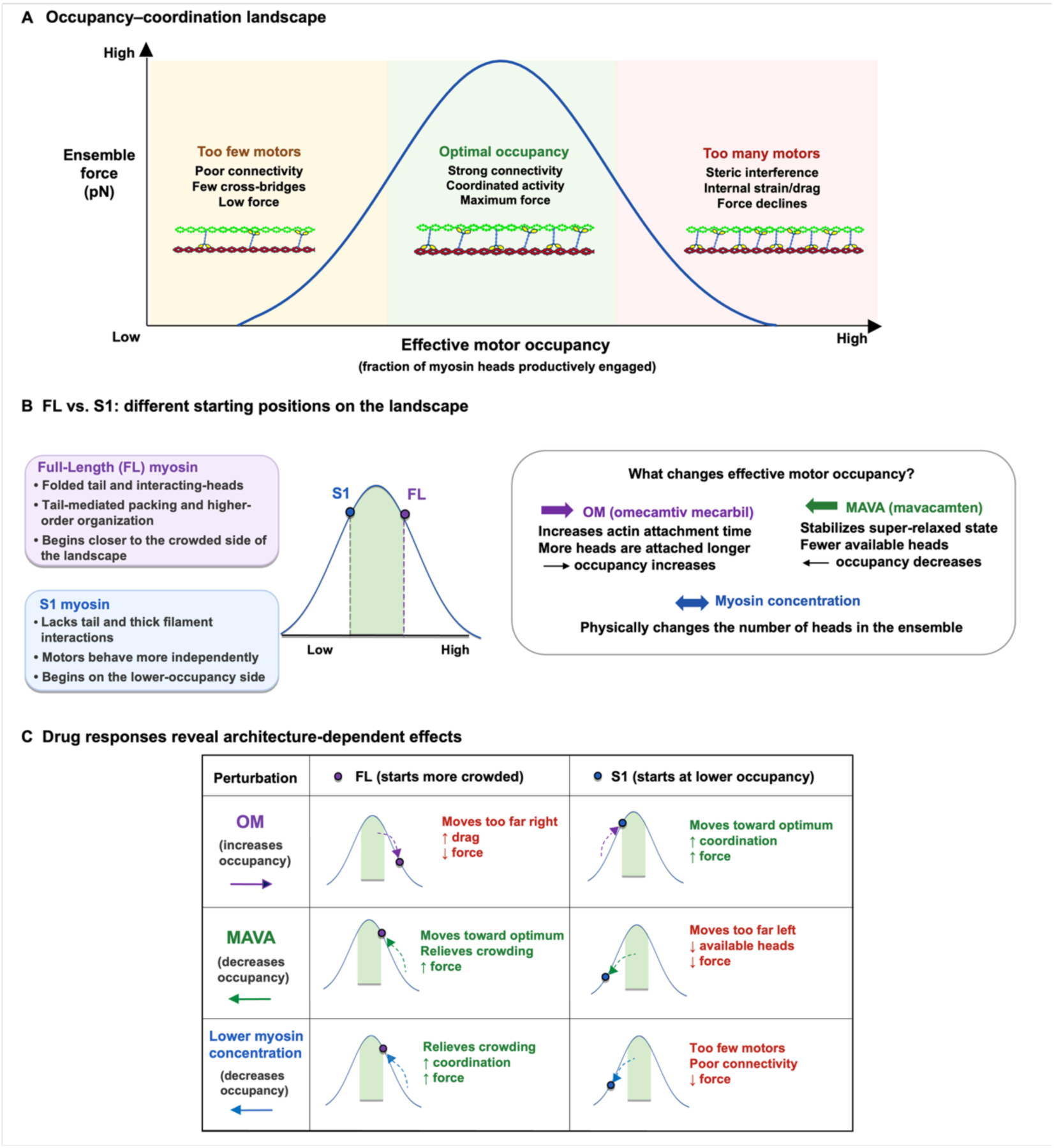
Proposed optimal motor-occupancy framework for architecture- and drug-dependent cardiac myosin ensemble force generation. **(A)** Conceptual relationship between effective motor occupancy and ensemble force. At low occupancy, too few engaged motors result in poor actin–myosin connectivity, limited cross-bridge formation, and low force. At an intermediate optimal occupancy, strong connectivity and coordinated motor activity produce maximal force. At excessive occupancy, steric interference, internal strain, and drag between motors reduce force output. **(B)** Proposed starting positions of full-length (FL) and S1 myosin ensembles along the occupancy landscape. FL myosin is proposed to occupy a different region of the landscape because its tail and two-headed architecture permit additional head–head, head–tail, and higher-order structural interactions that are absent from S1. S1 myosin, which lacks the tail and associated higher-order interactions, is proposed to begin on the lower-occupancy side and behave more independently. Omecamtiv mecarbil OM is proposed to increase effective motor engagement by prolonging actin-attachment time, whereas mavacamten (MAVA) suppresses cardiac myosin motor activity and can additionally reduce head availability through stabilization of autoinhibited/SRX states in two-headed myosin. Changing myosin solution concentration alters the number of motors available within the ensemble. **(C)** Schematic interpretation of architecture-dependent responses to experimental perturbations. Increasing occupancy with OM may shift FL ensembles beyond the optimal range, increasing interference and drag, while shifting S1 ensembles toward the optimal range. Decreasing effective motor engagement with MAVA may relieve counterproductive interactions at high loading while suppressing productive motor activity when initial loading is lower. Reducing myosin concentration may similarly relieve crowding at high initial occupancy or impair connectivity when initial occupancy is already low. The green region represents the proposed occupancy range that maximizes motor coordination and force generation. This framework represents a mechanistic interpretation of the force measurements; effective motor occupancy was not measured directly. FL, full-length; S1, myosin subfragment 1; OM, omecamtiv mecarbil; MAVA, mavacamten.

The S1 results provide the strongest statistical support for this loading-dependent interpretation. Both OM and MAVA produced significant myosin concentration × drug concentration interactions, demonstrating that the effect of each perturbation on maximum force observed in each trace depended on the initial myosin loading condition. These interactions are particularly informative because OM and MAVA alter motor engagement through distinct molecular mechanisms. OM prolongs actin attachment, whereas MAVA directly suppresses cardiac myosin motor activity (46, 49, 50, 53, 58). Despite these opposing molecular actions, the collective consequences of both perturbations depended on the starting state of the ensemble. These findings indicate that neither increasing nor decreasing effective motor engagement produces a uniform effect on collective force generation.

At higher S1 loading, OM increased mean maximum forces recorded in the traces at both drug concentrations, whereas MAVA produced an increase at the intermediate concentration followed by a decrease at the higher concentration. At lower S1 loading, OM reduced mean maximum force at 0.4 mg/mL before returning near the no-drug mean at 0.8 mg/mL, whereas MAVA produced a progressive decrease across increasing drug concentrations. Although these mean response patterns differed between the two compounds, the significant loading × drug interactions demonstrate that the effect of each pharmacological perturbation depended on the initial myosin-loading condition. These responses should also be considered in the context of the drug concentrations used. The OM and MAVA concentrations used may approach saturation conditions according to previous studies (54, 63, 78). A broader concentration series, including lower drug concentrations, will help clarify how pharmacological perturbations influence collective force generation in this system.

The FL ensembles exhibited a more complex response. Although the mean maximum-force responses to both OM and MAVA differed across loading conditions, neither loading × drug interaction reached statistical significance when maximum force generated within that bundle trace was used as the primary endpoint. These results therefore do not establish statistically significant loading-dependent drug responses in FL ensembles. However, the distinct mean-response patterns observed in FL ensembles may provide hypotheses regarding additional contributions from FL- specific structural features. Unlike S1, FL myosin retains the extended tail domain, introducing additional opportunities for molecular organization, steric hindrance, and mechanical interaction within the ensemble. Previous in vitro motility work with skeletal myosin has shown that the myosin tail can contribute to drag and reduce actin-filament velocity (43). In addition, FL myosin retains structural elements capable of head–head, head–tail, and higher-order interactions that are absent from the single-headed S1 construct (77, 79). Consequently, collective force generation in FL ensembles may reflect not only motor-domain activity and effective engagement but also tail-mediated resistance and higher-order ensemble organization.

This additional complexity may obscure relationships that are more readily resolved in the simplified S1 system. In S1 ensembles, changes in motor loading or drug-induced motor engagement may be reflected relatively directly in maximum force reached in each trace. In contrast, FL perturbations may simultaneously influence productive force generation, resistive interactions, molecular organization, and collective dynamics. The resulting effects may therefore be distributed across multiple features of ensemble behavior rather than appearing as consistent differences in maximum force alone. The comparison between FL and S1 thus raises the possibility that motor architecture influences how changes in motor engagement are translated into collective mechanical output, although the nonsignificant FL interactions do not establish a distinct architecture-dependent drug response.

The representative force trajectories provide supporting, but qualitative, evidence that maximum force recorded within a bundle trace does not capture all aspects of ensemble behavior. Across several representative conditions, changes in myosin loading or pharmacological perturbation were accompanied by visible differences in the magnitude and character of short-timescale force fluctuations. These observations did not follow a simple relationship with either motor concentration or drug treatment. For example, perturbations that altered effective motor engagement could increase force fluctuations under one loading condition while reducing them under another. Some trajectories continued to rise without reaching a clear plateau, indicating that maximum measured force should not be interpreted as a conventional stall force or as the intrinsic maximum force-generating capacity of the ensemble. The measured response may also reflect time-dependent changes in motor engagement, filament movement, or assembly organization. Consistent with this possibility, theoretical simulations of small myosin ensembles under elastic loading have shown that force-dependent motor attachment and detachment, including catch-bond behavior, can contribute to dynamic force responses (24). Although these simulations do not establish the mechanism underlying the present trajectories, they illustrate how motor kinetics and mechanical constraints can shape ensemble force development over time.

The mechanical environment of the assay also provides key context for interpreting these results. Unlike ensemble assays in which myosins are anchored to a rigid surface like a bead and interacts with an actin filament (80, 81), our assay uses a compliant, self-assembled configuration in which a bead is attached to one actin filament and another filament is anchored to the surface (15, 17). Myosin molecules are not rigidly anchored within the assembly and can interact dynamically with the filaments as they progress through the ATPase cycle. The resulting mechanical responses may therefore reflect evolving motor–filament interactions and ensemble organization rather than a single, persistent crosslinking arrangement. The observed bead displacement establishes a mechanical response transmitted through the reconstituted assembly but does not identify a unique molecular linkage responsible for that response.

This distinction is particularly relevant when considering S1, which contains a single motor head and lacks the extended tail of full-length myosin. Although previous studies have reported that S1 can promote actin-filament bundling through interactions involving the motor domain and ELC, the molecular basis of interfilament coupling in the present assay remains unresolved. Filament polarity was also not determined, leaving open the possibility that motors interacting with filaments of different orientations contribute mechanical effects that reinforce or oppose one another. However, unlike assays in which motors are rigidly anchored, this configuration allows force generation to emerge within a compliant assembly in which motor engagement and mechanical interactions evolve together. Although these coupled dynamics complicate the attribution of observed responses to specific molecular mechanisms, they provide an opportunity to investigate how force generation and mechanical feedback emerge from the interplay between motor activity and the evolving mechanical environment.

Collectively, the results support a model in which collective force generation emerges from the balance between motor engagement and mechanical coordination rather than motor number alone. Increasing the number of motors or prolonging their attachment can increase the potential for force generation, but these changes may also increase competition for interaction space, steric interference, internal strain, or resistive interactions. Conversely, reducing the number of available motors may either impair force generation or relieve mechanically counterproductive interactions, depending on the initial ensemble state. The consequences of changing effective motor occupancy therefore depend on where an ensemble begins within this balance.

This interpretation also provides a potential connection to previous work demonstrating mechanical coupling and load-dependent feedback within collective motor systems (8, 10, 11, 17, 19, 20). As motors attach to actin and generate force, they alter the mechanical environment experienced by other motors. Changes in motor number, attachment lifetime, or molecular architecture may therefore influence not only the contribution of individual motors but also the forces and constraints experienced by neighboring motors. Collective force generation can consequently emerge from feedback between motor kinetics, motor occupancy, actin mechanics, and ensemble organization.

The proposed occupancy landscape in **Figure 7** provides a conceptual model for integrating these observations. Ensembles with different initial myosin loading conditions may occupy different regions of a mechanical landscape in which productive force generation is limited at one extreme by insufficient motor engagement and at the other by excessive occupancy or mechanical interference. OM and MAVA shift the effective ensemble state in opposite directions, but the outcome of either perturbation depends on the initial position of the ensemble. This model does not imply that effective motor occupancy was measured directly or that specific occupancy states can be assigned to individual experimental conditions. Instead, it provides a physical interpretation for understanding why increasing or decreasing motor engagement can produce different collective outcomes depending on the starting ensemble state.

Future studies should examine a broader range of myosin-loading and drug concentrations to better define how these variables influence collective force generation. Accounting for filament polarity and organization, alongside further characterization of the interactions that couple filaments within the assembly, will help clarify how motor activity is translated into the measured mechanical response. Additional quantitative analysis of force-development kinetics, fluctuations, and persistence may also reveal differences in ensemble behavior that are not captured by maximum force alone.

### Conclusion

Together, the results are consistent with a model in which the mechanical output of reconstituted actomyosin assemblies depends on motor engagement and collective mechanical organization rather than myosin solution concentration alone. Using OM and MAVA as complementary perturbations of effective motor occupancy, we demonstrate that increasing or decreasing the number of engaged motors can either enhance or diminish collective force generation depending on the initial state of the ensemble. This reconciles the seemingly paradoxical observation that both prolonging actin attachment with OM and reducing the number of available myosin heads with MAVA can improve force under specific ensemble conditions.

More broadly, these results support the growing view that actomyosin assemblies exhibit emergent mechanical behavior that cannot be predicted solely from the properties of individual motors. Building on our previous observations in skeletal myosin ensembles, we propose that the actin filament serves as a mechanically responsive feedback element that integrates the actions of many individual motors into coordinated ensemble behavior. As such, ensemble occupancy modifies the mechanical state of actin, while the evolving mechanical state of actin feeds back to regulate motor activity.

This bidirectional mechanical coupling provides a conceptual mechanism by which ensembles can collectively sense and adapt to changes in motor occupancy without requiring direct communication between individual myosin molecules. Understanding these feedback mechanisms may ultimately provide new insights into how molecular regulation translates into coordinated contractility in both healthy and diseased muscle.

## Data Availability Statement

The data supporting the findings of this study are provided in the main text. Data are available upon reasonable request.

## Author Contributions

OYA and MAHC are co-first authors and were involved in all aspects of the work, including experiments, data analysis, and manuscript preparation. OYA, MAHC, HMS, and DNR aided in assay development, data acquisition, and analysis. OYA, MAHC, and DNR designed the experiments, analyzed data, and prepared the manuscript.

## Acknowledgements

This work was supported by American Heart Association #848586, National Institutes of Health R35GM147050, National Science Foundation CAREER #2439729, and National Science Foundation REU Site: Ole Miss Nanoengineering Summer REU Program #2148764.

## References

1. L. M. Coluccio, Myosins: A Superfamily of Molecular Motors (Springer Netherlands, 2008).

2. J. T. Finer, R. M. Simmons, J. A. Spudich, Single myosin molecule mechanics: piconewton forces and nanometre steps. Nature 368, 113–119 (1994).

3. J. T. Finer, A. D. Mehta, J. A. Spudich, Characterization of single actin-myosin interactions. Biophys J 68, 291S–296S; discussion 296S-297S (1995).

4. C. Rüegg, et al., Molecular motors: force and movement generated by single myosin II molecules. News Physiol Sci 17, 213–218 (2002).

5. R. W. Lymn, E. W. Taylor, Mechanism of adenosine triphosphate hydrolysis by actomyosin. Biochemistry 10, 4617–4624 (1971).

6. H. Huxley, J. Hanson, Changes in the Cross-Striations of Muscle during Contraction and Stretch and their Structural Interpretation. Nature 173, 973–976 (1954).

7. J. Robert-Paganin, O. Pylypenko, C. Kikuti, H. L. Sweeney, A. Houdusse, Force Generation by Myosin Motors: A Structural Perspective. Chem. Rev. 120, 5–35 (2020).

8. M. Kaya, Y. Tani, T. Washio, T. Hisada, H. Higuchi, Coordinated force generation of skeletal myosins in myofilaments through motor coupling. Nat Commun 8, 16036 (2017).

9. M. Kaya, H. Higuchi, Nonlinear elasticity and an 8-nm working stroke of single myosin molecules in myofilaments. Science 329, 686–689 (2010).

10. S. Walcott, D. M. Warshaw, E. P. Debold, Mechanical coupling between myosin molecules causes differences between ensemble and single-molecule measurements. Biophys J 103, 501–510 (2012).

11. L. Hilbert, S. Cumarasamy, N. B. Zitouni, M. C. Mackey, A.-M. Lauzon, The kinetics of mechanically coupled myosins exhibit group size-dependent regimes. Biophys J 105, 1466–1474 (2013).

12. J. A. Wagoner, K. A. Dill, Evolution of mechanical cooperativity among myosin II motors. Proceedings of the National Academy of Sciences 118, e2101871118 (2021).

13. T. J. Stewart, V. Murthy, S. P. Dugan, J. E. Baker, Velocity of myosin-based actin sliding depends on attachment and detachment kinetics and reaches a maximum when myosin-binding sites on actin saturate. J Biol Chem 297, 101178 (2021).

14. O. Al Azzam, C. L. Trussell, D. N. Reinemann, Measuring force generation within reconstituted microtubule bundle assemblies using optical tweezers. Cytoskeleton (Hoboken) 78, 111–125 (2021).

15. O. Al Azzam, J. C. Watts, J. E. Reynolds, J. E. Davis, D. N. Reinemann, Probing Myosin Ensemble Mechanics in Actin Filament Bundles Using Optical Tweezers. JoVE 63672 (2022). 10.3791/63672.

16. M. J. Greenberg, J. R. Moore, The Molecular Basis of Frictional Loads in the In Vitro Motility Assay with Applications to the Study of the Loaded Mechanochemistry of Molecular Motors. Cytoskeleton (Hoboken) 67, 273–285 (2010).

17. O. Y. Al Azzam, J. C. Watts, J. E. Reynolds, J. E. Davis, D. N. Reinemann, Myosin II Adjusts Motility Properties and Regulates Force Production Based on Motor Environment. Cell Mol Bioeng 15, 451–465 (2022).

18. S. K. Barrick, M. J. Greenberg, Cardiac myosin contraction and mechanotransduction in health and disease. J Biol Chem 297, 101297 (2021).

19. M. J. Greenberg, H. Shuman, E. M. Ostap, Inherent force-dependent properties of β-cardiac myosin contribute to the force-velocity relationship of cardiac muscle. Biophys J 107, L41–L44 (2014).

20. J. Sung, et al., Harmonic force spectroscopy measures load-dependent kinetics of individual human β-cardiac myosin molecules. Nat Commun 6, 7931 (2015).

21. M. Caremani, L. Melli, M. Dolfi, V. Lombardi, M. Linari, Force and number of myosin motors during muscle shortening and the coupling with the release of the ATP hydrolysis products. J Physiol 593, 3313–3332 (2015).

22. D. R. Jackson, J. E. Baker, The Energetics of Allosteric Regulation of ADP Release From Myosin Heads. Phys Chem Chem Phys 11, 4808–4814 (2009).

23. M. Kovács, K. Thirumurugan, P. J. Knight, J. R. Sellers, Load-dependent mechanism of nonmuscle myosin 2. Proc Natl Acad Sci U S A 104, 9994–9999 (2007).

24. T. Erdmann, P. J. Albert, U. S. Schwarz, Stochastic dynamics of small ensembles of non-processive molecular motors: the parallel cluster model. J Chem Phys 139, 175104 (2013).

25. V. N. Amari, E. M. Kerivan, D. N. Reinemann, Measuring emergent mechanical changes in cytoskeletal ensembles in vitro using QCM-D. Front. Cell Dev. Biol. 13 (2025).

26. C. Veigel, J. E. Molloy, S. Schmitz, J. Kendrick-Jones, Load-dependent kinetics of force production by smooth muscle myosin measured with optical tweezers. Nat Cell Biol 5, 980–986 (2003).

27. S. M. Mijailovich, et al., Modeling the Actin.myosin ATPase Cross-Bridge Cycle for Skeletal and Cardiac Muscle Myosin Isoforms. Biophys J 112, 984–996 (2017).

28. J. Grewe, U. S. Schwarz, Mechanosensitive self-assembly of myosin II minifilaments. Phys Rev E 101, 022402 (2020).

29. A. P. Baldo, J. C. Tardiff, S. D. Schwartz, Mechanochemical Function of Myosin II: Investigation into the Recovery Stroke and ATP Hydrolysis. J Phys Chem B 124, 10014–10023 (2020).

30. D. E. Rassier, A. Månsson, Mechanisms of myosin II force generation: insights from novel experimental techniques and approaches. Physiol Rev 105, 1–93 (2025).

31. B. Alberts, et al., Molecular Biology of the Cell, 4th Ed. (Garland Science, 2002).

32. S. K. Gollapudi, M. Yu, Q.-F. Gan, S. Nag, Synthetic thick filaments: A new avenue for better understanding the myosin super-relaxed state in healthy, diseased, and mavacamten-treated cardiac systems. J Biol Chem 296, 100114 (2020).

33. M. J. Bloemink, M. A. Geeves, Shaking the Myosin Family Tree Biochemical kinetics defines four types of myosin motor. Semin Cell Dev Biol 22, 961–967 (2011).

34. J. Walklate, Z. Ujfalusi, M. A. Geeves, Myosin isoforms and the mechanochemical cross-bridge cycle. J Exp Biol 219, 168–174 (2016).

35. A. M. Gordon, M. Regnier, E. Homsher, Skeletal and Cardiac Muscle Contractile Activation: Tropomyosin “Rocks and Rolls.” Physiology 16, 49–55 (2001).

36. A. Nayak, et al., Single-molecule analysis reveals that regulatory light chains fine-tune skeletal myosin II function. J Biol Chem 295, 7046–7059 (2020).

37. S. M. Day, J. C. Tardiff, E. M. Ostap, Myosin modulators: emerging approaches for the treatment of cardiomyopathies and heart failure. J Clin Invest 132 (2022).

38. M. Kawana, J. A. Spudich, K. M. Ruppel, Hypertrophic cardiomyopathy: Mutations to mechanisms to therapies. Front Physiol 13, 975076 (2022).

39. J. A. Spudich, N. Nandwani, J. Robert-Paganin, A. Houdusse, K. M. Ruppel, Reassessing the unifying hypothesis for hypercontractility caused by myosin mutations in hypertrophic cardiomyopathy. EMBO J 43, 4139–4155 (2024).

40. J. C. Deacon, M. J. Bloemink, H. Rezavandi, M. A. Geeves, L. A. Leinwand, Identification of functional differences between recombinant human α and β cardiac myosin motors. Cell Mol Life Sci 69, 2261–2277 (2012).

41. C. Liu, M. Kawana, D. Song, K. M. Ruppel, J. A. Spudich, Controlling load-dependent kinetics of β-cardiac myosin at the single molecule level. Nat Struct Mol Biol 25, 505–514 (2018).

42. T. Wang, A. Nayak, T. Kraft, M. Amrute-Nayak, Single-Molecule Investigation of Load-Dependent Actomyosin Dissociation Kinetics for Cardiac and Slow Skeletal Myosin. Small 20, e2406865 (2024).

43. B. Guo, W. H. Guilford, The tail of myosin reduces actin filament velocity in the in vitro motility assay. Cell Motil Cytoskeleton 59, 264–272 (2004).

44. T. Ando, D. Scales, Skeletal muscle myosin subfragment-1 induces bundle formation by actin filaments. J Biol Chem 260, 2321–2327 (1985).

45. T. Miyanishi, T. Ishikawa, T. Hayashibara, T. Maita, T. Wakabayashi, The two actin-binding regions on the myosin heads of cardiac muscle. Biochemistry 41, 5429–5438 (2002).

46. K. Kazmierczak, et al., The role of the N-terminus of the myosin essential light chain in cardiac muscle contraction. J Mol Biol 387, 706–725 (2009).

47. D. J. Timson, Fine tuning the myosin motor: the role of the essential light chain in striated muscle myosin. Biochimie 85, 639–645 (2003).

48. M. S. Miller, et al., The essential light chain N-terminal extension alters force and fiber kinetics in mouse cardiac muscle. J Biol Chem 280, 34427–34434 (2005).

49. S. Lowey, L. D. Saraswat, H. Liu, N. Volkmann, D. Hanein, Evidence for an Interaction between the SH3 Domain and the N-terminal Extension of the Essential Light Chain in Class II Myosins. J Mol Biol 371, 902–913 (2007).

50. P. Muthu, et al., Structural and functional aspects of the myosin essential light chain in cardiac muscle contraction. FASEB J 25, 4394–4405 (2011).

51. E. M. Aydt, G. Wolff, I. Morano, Molecular modeling of the myosin-S1(A1) isoform. J Struct Biol 159, 158–163 (2007).

52. Y. H. Sitbon, S. Yadav, K. Kazmierczak, D. S. Cordary, Insights into myosin regulatory and essential light chains: a focus on their roles in cardiac and skeletal muscle function, development and disease. J Muscle Res Cell Motil 41, 313–327 (2020).

53. D. Szczesna-Cordary, Modulating the myosin super-relaxed state as a therapeutic strategy for myosin light chain–linked cardiomyopathies. Biophys Rev 18, 641–648 (2026).

54. M. S. Woody, et al., Positive cardiac inotrope omecamtiv mecarbil activates muscle despite suppressing the myosin working stroke. Nat Commun 9, 3838 (2018).

55. E. M. Green, et al., A small-molecule inhibitor of sarcomere contractility suppresses hypertrophic cardiomyopathy in mice. Science 351, 617–621 (2016).

56. T. Aksel, E. Choe Yu, S. Sutton, K. M. Ruppel, J. A. Spudich, Ensemble force changes that result from human cardiac myosin mutations and a small-molecule effector. Cell Rep 11, 910–920 (2015).

57. M. J. Daniels, L. Fusi, C. Semsarian, S. S. Naidu, Myosin Modulation in Hypertrophic Cardiomyopathy and Systolic Heart Failure: Getting Inside the Engine. Circulation 144, 759–762 (2021).

58. F. I. Malik, et al., Cardiac myosin activation: a potential therapeutic approach for systolic heart failure. Science 331, 1439–1443 (2011).

59. D. Auguin, et al., Omecamtiv mecarbil and Mavacamten target the same myosin pocket despite opposite effects in heart contraction. Nat Commun 15, 4885 (2024).

60. K. R. L. Reyes, G. Bilgili, F. Rader, Mavacamten: A First-in-class Oral Modulator of Cardiac Myosin for the Treatment of Symptomatic Hypertrophic Obstructive Cardiomyopathy. Heart Int 16, 91–98 (2022).

61. A. Tower-Rader, J. Ramchand, S. E. Nissen, M. Y. Desai, Mavacamten: a novel small molecule modulator of β-cardiac myosin for treatment of hypertrophic cardiomyopathy. Expert Opinion on Investigational Drugs 29, 1171–1178 (2020).

62. V. J. Planelles-Herrero, J. J. Hartman, J. Robert-Paganin, F. I. Malik, A. Houdusse, Mechanistic and structural basis for activation of cardiac myosin force production by omecamtiv mecarbil. Nat Commun 8, 190 (2017).

63. A. M. Swenson, et al., Omecamtiv Mecarbil Enhances the Duty Ratio of Human β-Cardiac Myosin Resulting in Increased Calcium Sensitivity and Slowed Force Development in Cardiac Muscle. J Biol Chem 292, 3768–3778 (2017).

64. J. R. Teerlink, et al., Dose-dependent augmentation of cardiac systolic function with the selective cardiac myosin activator, omecamtiv mecarbil: a first-in-man study. Lancet 378, 667–675 (2011).

65. J. R. Teerlink, et al., Cardiac Myosin Activation with Omecamtiv Mecarbil in Systolic Heart Failure. New England Journal of Medicine 384, 105–116 (2021).

66. R. L. Anderson, et al., Deciphering the super relaxed state of human β-cardiac myosin and the mode of action of mavacamten from myosin molecules to muscle fibers. Proc Natl Acad Sci U S A 115, E8143–E8152 (2018).

67. T. Dong, B. Alencherry, S. Ospina, M. Y. Desai, Review of Mavacamten for Obstructive Hypertrophic Cardiomyopathy and Future Directions. Drug Des Devel Ther 17, 1097–1106 (2023).

68. S. Nag, S. K. Gollapudi, C. L. del Rio, J. A. Spudich, R. McDowell, Mavacamten, a precision medicine for hypertrophic cardiomyopathy: From a motor protein to patients. Science Advances 9, eabo7622 (2023).

69. I. Olivotto, et al., Mavacamten for treatment of symptomatic obstructive hypertrophic cardiomyopathy (EXPLORER-HCM): a randomised, double-blind, placebo-controlled, phase 3 trial. The Lancet 396, 759–769 (2020).

70. S. N. McMillan, J. R. T. Pitts, B. Barua, D. A. Winkelmann, C. A. Scarff, Mavacamten inhibits myosin activity by stabilizing the myosin interacting-heads motif and stalling motor force generation. Sci Adv 12, eaea9335 (2026).

71. D. Auguin, et al., Omecamtiv mecarbil and Mavacamten target the same myosin pocket despite opposite effects in heart contraction. Nat Commun 15, 4885 (2024).

72. M. P. Grillo, et al., In vitro and in vivo pharmacokinetic characterization of mavacamten, a first-in-class small molecule allosteric modulator of beta cardiac myosin. Xenobiotica 49, 718–733 (2019).

73. E. Spahiu, E. Kastrati, M. Amrute-Nayak, PyChelator: a Python-based Colab and web application for metal chelator calculations. BMC Bioinformatics 25, 239 (2024).

74. D. E. Bentil, How Brownian displacement goes down during actin-myosin interaction in the laser trap. Mathematical and Computer Modelling 29, 131–144 (1999).

75. M. Capitanio, F. S. Pavone, Interrogating Biology with Force: Single Molecule High-Resolution Measurements with Optical Tweezers. Biophys J 105, 1293–1303 (2013).

76. C. Barutel, S. Fürthauer, Entropic, active, and frictional forces in cytoskeletal crosslinkingPhys. Rev. Res. 8, 033122 (2026).

77. J. A. Rohde, O. Roopnarine, D. D. Thomas, J. M. Muretta, Mavacamten stabilizes an autoinhibited state of two-headed cardiac myosin. Proc Natl Acad Sci U S A 115, E7486–E7494 (2018).

78. R. F. Kawas, et al., A small-molecule modulator of cardiac myosin acts on multiple stages of the myosin chemomechanical cycle. J Biol Chem 292, 16571–16577 (2017).

79. R. Craig, R. Padrón, Structural basis of the super-and hyper-relaxed states of myosin II. J Gen Physiol 154, e202113012 (2022).

80. E. P. Debold, J. B. Patlak, D. M. Warshaw, Slip Sliding Away: Load-Dependence of Velocity Generated by Skeletal Muscle Myosin Molecules in the Laser Trap. Biophys J 89, L34–L36 (2005).

81. I. Pertici, et al., A myosin II nanomachine mimicking the striated muscle. Nat Commun 9, 3532 (2018).

